# A transient signal in foveal superior colliculus neurons when jumpstarting peripheral saccadic orienting

**DOI:** 10.64898/2026.04.06.716792

**Authors:** Tong Zhang, Anna F. Denninger, Ziad M. Hafed

**Affiliations:** Werner Reichardt Centre for Integrative Neuroscience, University of Tübingen, Tübingen, Germany; Hertie Institute for Clinical Brain Research, University of Tübingen, Tübingen, Germany; Department of Psychiatry and Psychotherapy, Center for Mental Health (TüCMH), University of Tübingen, Tübingen, Germany

**Keywords:** Saccades, superior colliculus, primary visual cortex, delayed saccade task, fovea

## Abstract

The superior colliculus (SC) both senses the environment and orients gaze within it. While the SC’s sensory and motor bursts appear qualitatively similar to each other, population activity structure in the two processing regimes (visual and motor) is very different, necessitating a hitherto unexplored rapid representational transformation, occurring on the scale of only tens of milliseconds. Here, we first show that when a planned saccade is released with a go signal, peripheral SC neurons representing the saccade target location exhibit a transient, short-latency pause right before their motor bursts eventually emerge. This pause starts within ∼50 ms from the go signal; it is stimulus-dependent; and it is significantly weaker in purely visual SC neurons than in saccade-related ones. Foveal SC neurons, on the other hand, transiently burst, and their bursts lead the peripheral neurons’ pauses by ∼10 milliseconds. These transient foveal bursts also lead the same foveal neurons’ saccade-related pauses, which are time-locked to movement generation. Remarkably, during immediate visually-guided saccade tasks, requiring a transformation from visual to motor peripheral bursts in <50-100 ms, the transient foveal SC bursts still occur, resulting in simultaneous short-latency bursting at two disparate SC loci: one foveal; and one eccentric and responding to the visual appearance of the saccade target. Our results suggest that in classic saccade tasks used to investigate visual, motor, and cognitive processes in primate brains, a transient foveal SC signal precedes peripheral saccadic orienting, and may facilitate a necessary rapid representational transformation needed for SC saccade motor bursts to ensue.

**Significance:** Saccade control studies often involve behavioral paradigms dissociating stimulus presentation, deliberation epochs, and “go” instructions. While the stimulus presentation, deliberation, and actual movement aspects have been well studied, the “go” phases of trials are less considered. We show that foveal superior colliculus neurons exhibit transient activity bursts at the “go” instruction for saccades, likely enabling a rapid representational transformations from a visual to a motor regime in the extrafoveal collicular neurons driving the eye movements.

## Introduction

Active vision entails a continual cycling between sensing, deliberation, and action (1–4). In the oculomotor system, the neuronal mechanisms underlying the different subcomponents of active vision have classically been studied using behavioral paradigms that attempt to dissociate these subcomponents from one another as much as possible. For example, in the delayed visually-guided saccade paradigm (5, 6), a visual stimulus first appears, but the initially fixated spot remains visible. This prevents a reflexive saccade to the appearing stimulus and allows investigating visual sensory processing mechanisms independent of an overt motor output. Then, after some delay, during which cognitive processes related to deliberation and motor planning may be investigated, the initially fixated spot disappears, allowing a saccade towards the eccentric stimulus to be subsequently triggered. In this case, motor-related processes associated with saccade generation can be studied under a constant, steady-state visual appearance of the environment.

Among the many insights gleaned from the classic delayed saccade paradigm, it was recently recognized that this paradigm additionally highlights a fundamental problem in sensory-driven motor behavior, namely the need to transform neuronal representations from a sensory regime to a motor regime, often in the very same neurons and within very short time intervals (7–14). For example, population activity subspaces in the superior colliculus (SC) are different from each other at stimulus onset and saccade generation (7, 8). And, at the individual neuron level, some SC neurons can prefer a particular visual image feature in the stimulus onset phase of trials, but a different image feature in the saccade generation phase (8). This rapid alteration in operating regimes of identical neurons likely necessitates a switch-like mechanism at some point during delayed saccade task trials, and this is what we investigated here.

We specifically hypothesized that the go, or release, signal for saccades may be associated with neuronal dynamics that were not previously characterized in sufficient detail. Indeed, past studies generally focused on neuronal activity for (peripheral) neurons representing the eccentric locations of the saccade targets. These studies uncovered well known sensory, cognitive, and motor processes associated with the neurons driving the eye movements. However, it was not known what specifically happens to foveal representations in the SC at the time of the go signal: since active vision ultimately starts and ends with the fovea, understanding foveal SC representations when releasing saccades is important.

To address this gap, we studied the activity of peripheral SC and primary visual cortex (V1) neurons during the delayed saccade paradigm, with a particular focus on neuronal activity around the time of the go signal. Specifically, we looked at the release instruction phase of the task, unlike prior work which has predominantly focused on visual responses or on discharges associated with the actual initiation of the saccades themselves. We found that releasing an instructed eye movement in this delayed saccade paradigm is associated with a short-latency transient pause in SC, but not V1, activity right before SC saccade-related motor bursts ensue. Importantly, this pause is preceded by foveal SC bursts ∼10 ms earlier, and the foveal burst properties suggest that they could be a generalized trigger signal rather than a stimulus-dependent phenomenon. These foveal bursts explicitly occur at a distinct time from previously studied foveal SC neuronal modulations as a result of either peripheral stimulus onset or actual saccade execution (15–18). Remarkably, when we switched our paradigm to instead employ immediate visually-guided saccades, SC foveal bursts still happened, and thus temporally coincided with peripheral visual responses to target onsets in other SC neurons.

Our results reveal novel neuronal dynamics in the foveal SC at the time of the go signal in classic eye movement paradigms that have been used for several decades to study perception, cognition, and action. These results also motivate investigating mechanisms of foveal-to-peripheral, and vice-versa, neuronal modulations in the SC and other brain structures, especially when these modulations may not always be trivially explained by classic lateral inhibitory mechanisms (19–24).

## Materials and methods

### Research animals and ethical approvals

This study involved a re-analysis of data collected previously for other publications investigating other research questions. In all cases, we analyzed data collected from male rhesus macaque monkeys. For analyzing peripheral SC and V1 activity in the delayed visually-guided saccade task (see below for the details of the different behavioral tasks), we used the same database as the one in (8). For analyzing foveal SC activity in the delayed visually-guided saccade task, we used the control conditions of (15). And, for analyzing peripheral and foveal SC activity in the immediate visually-guided saccade task, we employed the control conditions of (25).

In all cases, all experiments were approved by ethics committees at the regional governmental offices of Tübingen.

### Behavioral tasks

#### Saccades-to-gratings task for peripheral SC and V1 neurons

For peripheral SC and V1 activity in the delayed saccade condition, the behavioral task was the “saccades-to-gratings” paradigm (8). This task was a slight modification of the classic delayed visually-guided saccade paradigm. Briefly, the monkeys first saw a white fixation spot of 10.8 by 10.8 min arc dimensions and 79.9 cd/m^2^ luminance. The gray background had a luminance of 26.11 cd/m^2^. After they fixated the white spot by a few hundred milliseconds, a visual target appeared eccentrically, centered on the recorded neurons’ response field (RF) locations. After maintaining fixation for another few hundred milliseconds after target onset, the fixation spot was removed, and this was the cue for the monkeys to generate a visually-guided saccade towards the eccentric target. Thus, the fixation spot removal was the go signal in this task, as is the case in classic instantiations of the same paradigm.

The visual target in our task consisted of a disc of 3 deg radius, the inside of which had a sine wave grating. We always placed a white spot (like the fixation spot) at the center of the disc, surrounded by a small gray disc (to avoid visibility loss of the spot due to the background sine wave grating). This placement allowed the saccades to be accurately made towards the disc’s center, which was critical for the previous study (but it did not influence the scientific questions of the present one). In the spatial frequency version of the task, the sine wave grating was vertical and had 100% contrast, but it could have different spatial frequencies across different trials. In the contrast version of the task, we fixed the spatial frequency to 1 cycle/deg (1 cpd) and instead varied the contrast of the vertical grating from trial to trial. And, finally, in the orientation version of the task, both the spatial frequency (1 cpd) and contrast (100%) were fixed, and the orientation of the grating was varied from trial to trial. Trials from each version of the task were collected in separate blocks.

#### Saccades-to-gratings task for foveal SC neurons

We also tested foveal SC activity in a similar delayed saccade paradigm; this allowed us to explore foveal modulations associated with the go signal epochs of the behavioral trials (see Results). In this case, the behavioral task was similar to the one above (15). Specifically, after fixating the fixation spot, an eccentric target appeared. Again, it consisted of a disc of 3 deg radius, the inside of which had a vertical sine wave grating of 100% contrast and either 1 cpd (low) or 4 cpd (high) spatial frequency. There was no spot placed at the center of the grating in this case. The target appeared at 8 deg eccentricity either to the right or left of fixation (except in two sessions in which it was placed at an oblique position of similar eccentricity; ∼10 deg). After maintaining fixation for a few hundred milliseconds after target onset, the fixation spot was removed, instructing the generation of the saccade towards the eccentric grating. After fixating the grating for another 500 ms, this grating was removed, and the monkeys were rewarded. A short inter-trial interval (with a blank gray screen) then ensued.

#### Immediate visually-guided saccade task for perihpheral and foveal SC neurons

For peripheral and foveal SC activity in the immediate saccade condition, we employed a classic reflexive visually-guided saccade paradigm; the control condition of (25). Specifically, the monkeys fixated a white fixation spot. After a few hundred milliseconds, the fixation spot was removed and a simultaneous saccade target (white disc of 0.51 deg radius) appeared either at the peripheral neurons’ RF positions or at an eccentricity >3.5 deg from the fovea for the foveal neurons.

#### RF mapping tasks

The delayed saccade tasks above allowed us to investigate neuronal dynamics at the time of transitioning from a visual regime (after stimulus onset and waiting for saccade instruction) to a motor regime (after the go signal). The reflexive saccade task, on the other hand, allowed us to explore what happens when the transition from visual to motor regimes needed to happen much more urgently, as quickly as possible after peripheral target onset. In all cases, we also mapped neuronal RF’s in the same sessions, and this was done in order to identify the RF’s positions relative to the eccentric saccade target. Our RF mapping tasks (26–29) involved fixating a fixation spot, like in all tasks above. After a few hundred milliseconds of fixation, a white spot (of the same dimensions as the fixation spot) appeared at some location on the display (pseudorandomly selected on every trial), and it typically remained on for 300-500 ms, before being turned off again. Across trials, we changed the position of the appearing spot, in order to construct the RF maps of the neurons.

### Data analysis and statistical tests

All saccadic eye movements were previously detected for the purposes of the earlier studies (8, 15, 25). Similarly, all neuronal preprocessing (such as spike sorting) was performed earlier. Our subsequent analyses were geared for the specific scientific purposes of the present study.

To characterize visual RF’s of neurons (e.g. Figs. 1A, C, 5A, C in Results), we averaged firing rate in an interval 50-150 ms after stimulus onset, and we plotted it as a function of the horizontal and vertical position of the appearing stimulus (including interpolation in between sampled positions). For some analyses, we also characterized the stimulus-offset responses of the recorded neurons from the RF mapping data (particularly for the foveal neurons in the delayed saccade paradigm). In this case, we simply measured average firing rate 50-150 ms after the offset of the target in the same RF mapping task. As we explain in Results, this analysis was aimed at checking whether the removal of the fixation spot in our main saccade paradigms caused sensory offset responses in the foveal neurons. Thus, for measuring such sensory offset responses from the RF mapping task, we picked stimulus locations in the RF mapping data that were consistent with the eccentricity of the fixation spot (in the main tasks) from the foveal neuron’s RF hotspot location (being defined as the point relative to gaze position for which the RF of the neuron exhibited the strongest visual response). In other words, because foveal SC neurons are strongly lateralized (29), fixation spot removal in the main tasks involved an offset of a stimulus that was at some non-zero eccentricity distance, *r*, from the hotspot of the recorded foveal neurons. Therefore, from the RF mapping data, we defined a ring around the hotspot location of each foveal SC neuron with radius of *r* deg (+/- 0.25 deg), where *r* was the eccentricity of the RF hotspot from the fixation spot. Then, we picked all sampled RF mapping trials in this ring of locations that were also in the contralateral hemifield relative to the recorded SC side (we picked contralateral locations to maximize the likelihood of observing an offset response if it did exist), and we measured sensory offset responses from these trials in particular. We then compared these sensory offset responses to those associated with the go signal in the main saccade tasks (which are described in Results).

**Figure 1.**
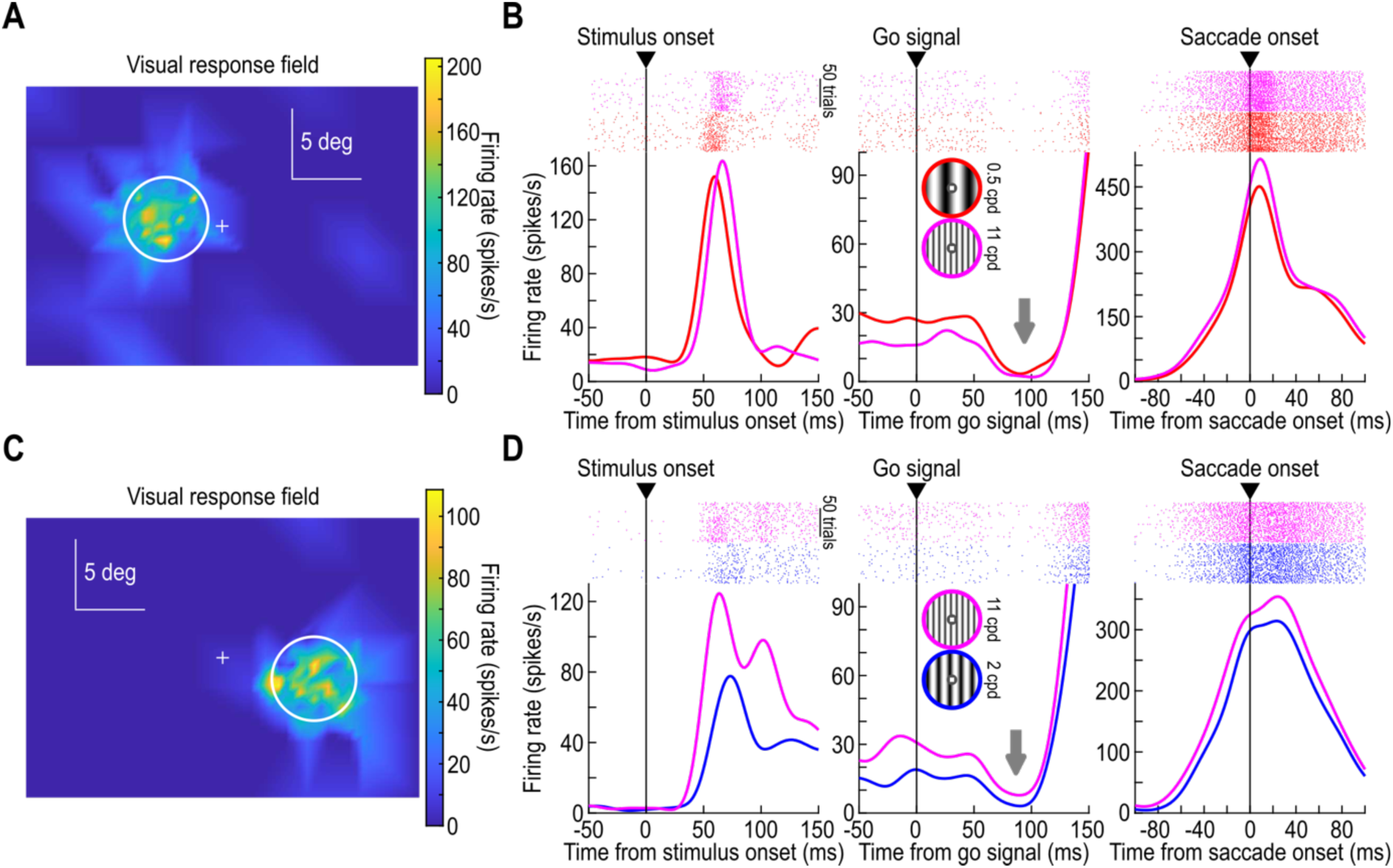
Transient activity pauses before the occurrence of saccade-related motor bursts in the SC. **(A)** Visual response field of an example SC neuron. The white circle indicates the saccade-target location and size; monkeys were only allowed to generate a saccade to the target upon a go signal (consisting of the removal of the fixation spot; Materials and methods) (8). **(B)** The activity of the same example neuron aligned to stimulus onset in the main saccade task (left), go signal onset (middle), and actual saccade onset (right). Two visual target appearances of the saccade target are shown (each target had a small central spot to aid in saccade accuracy) (8). Expected differences in visual (left) or motor (right) burst properties as a function of image appearance were observed (8, 28, 38, 39, 41, 42). Critically, shortly after the go signal (middle), the neuron paused its activity before the strong saccade-related burst could erupt. **(C, D)** Similar observations from a second example neuron, showing a time-locked activity pause after go signal before the motor bursts finally occurred (middle panel in **D**). Numbers of trial repetitions per panel can be seen from the individual trial spike rasters shown.

For the main saccade tasks, we plotted firing rates as a function of time from target onset, saccade onset, and go signal onset (fixation spot removal). The first two kinds of plots are similar to what was done previously, by us (8, 15, 25) and others; we included them here for providing the context associated with the third kind of plot, which was of interest for us in the current study. As we show in Results, this third plot gave either transient pauses or transient bursts in activity (shortly after the go signal), depending on the neuron location and behavioral task. To characterize these transient responses, we measured average firing rate in the interval 50-150 ms after go signal onset. For a reference, we also measured firing rate in the final 100 ms before go signal onset. This gave a baseline to which we compared go-signal responses when assessing whether there was a burst or pause occurring after the go signal. Such a comparison was made either by plotting the two measures directly against each other for each neuron, or by obtaining a neuronal modulation index. When assessing transient pauses (see Results), we defined the modulation index as the firing rate after the go signal (50-100 ms) minus the firing rate before the go signal (final 100 ms), divided by the sum of the two firing rates. Negative modulation indices meant reductions in firing rate after the go signal.

We also sometimes classified trial conditions (e.g. different spatial frequencies in the spatial frequency task) as a function of sustained activity level they gave in a given neuron at the end of the delay period (right before go signal onset). Specifically, we looked for the condition that gave the highest sustained activity and called it the most preferred sustained feature. Similarly, we looked for the image condition that gave the lowest sustained activity, and we called it the least preferred sustained feature. This allowed us to check whether transient pauses after the go signal that we characterize in Results depended on the starting firing rate that was present at the time of the go signal.

Similarly, we divided SC neurons according to their functional type (such as visual or visual-motor neurons). We used classifications that we used previously (8). Thus, we did not reclassify the neurons here.

To compare transient pauses and bursts directly (e.g. Fig. 7 in Results), we normalized individual neuron firing rates and then combined neurons by averaging their normalized firing rate curves. The normalization was achieved by measuring, for each neuron, the average firing rate in the final 100 ms before go signal onset, and then dividing the entire firing rate curve by this value. Thus, the normalized firing rate was 1 at the go signal, and bursts/pauses were higher/lower than 1. Sometimes, we also kept firing rates unnormalized, but we baseline-subtracted them based on the final 100 ms before the go signal. Thus, in this case, pauses were associated with negative baseline-subtracted firing rates.

Similarly, to assess the relative timing between transient pauses and transient bursts in the delayed saccade paradigm (e.g. Fig. 7 in Results), we relied on individual spike times. Across all trials and all neurons (either foveal or peripheral), we binned spike times around the go signal into 2-ms non-overlapping time bins. Then, we plotted a histogram of all spike times. For the peripheral neurons, they paused after the go signal (see Results). Thus, starting from 10 ms after the go signal, we searched for the first time point at which the histogram of spike times had 3 consecutive drops in spike likelihood after the go signal. This was considered the population latency of the pause. For the foveal neurons, which burst instead (see Results), we searched for the first time point at which the histogram of spike times had 3 consecutive increases in spike likelihood after the go signal. This was considered the population latency of the burst.

For foveal neurons, we also sometimes looked at task context. Specifically, we compared foveal bursts that we got from the go signal (see Results) to potential bursts that might be associated with the offset of a foveal target stimulating the recorded neurons’ RF’s. We obtained the latter from the ends of the trials in the delayed saccade paradigm used to study the foveal neurons. As mentioned above, after the monkeys fixated the grating with a saccade, the grating covered the fovea, and thus stimulated the recorded foveal neurons’ RF’s (15). When the grating disappeared, signaling the end of the trial, the monkeys were without any task-instruction and could look wherever they wanted. We measured firing rates after such grating disappearance, to look for potential stimulus-offset responses in the neurons. To ensure that there were no influences of potential saccades, we only included cases in which there were no saccades after grating disappearance for at least 150 ms. This way, we had an offset of a visual stimulus in the recorded RF’s, but there was no instructed saccade context (as was the case with the fixation spot offset of the go signal in the main task).

For the immediate saccade task, we used similar analyses to the ones above. For population firing rates, we normalized the firing rate of each neuron by the peak visual or foveal burst response (see Results). Specifically, we first subtracted the average activity of each neuron in the final 100 ms before stimulus onset from all firing rates at all times in the trials (25). Then, we divided the firing rate curves by the peak value occurring 50-100 ms after stimulus onset (25). Then, we averaged across all neurons. This allowed us to focus on the timing dynamics between foveal and peripheral bursts in this task, as well as on the relationships of the bursts to saccadic reaction times.

For such saccadic reaction time relationships, for each neuron, we split the trials into ones with saccadic reaction times faster than the median value of the session and ones with longer reaction times than the median value of the session. Then, we evaluated the peripheral and foveal bursts for both groups of trials separately across neurons.

Statistically, we always showed standard error of the mean ranges in all plots. For comparisons between conditions, we used signrank tests, rank sum tests, or t-tests. Sometimes, we split foveal SC neurons as ones with or without a foveal burst at the go signal. This classification was made statistically. If the neuron’s firing rate 50-150 ms after the go signal was significantly higher than in the final 100 ms before the go signal with an t-test, then the neuron was classified as having a foveal burst; otherwise, it was not. This allowed us to compare foveal bursts to stimulus-offset responses from RF mapping tasks.

## Results

We first investigated the dynamics of switching between fixation and saccade generation in the standard delayed saccade paradigm. In this paradigm, monkeys fixate an initial fixation spot, and an eccentric saccade target appears. The monkeys withhold saccadic orienting towards the eccentric target until a go signal arrives, which comes in the form of fixation spot disappearance. Classic SC recordings of peripheral neurons in this task demonstrate visual responses to target onset followed by motor bursts at saccade onset, often in the very same neurons (6–8, 30, 31). Here, we were interested in the neuronal dynamics that take place in between these two phases, and particularly at the time of the go signal. Moreover, from the same task, we also recorded from foveal SC neurons, which presumably should not burst for either target or saccade onset because of the disparity between their RF locations and the saccade target location (16–19, 32–37).

In what follows, we first start by characterizing peripheral SC and V1 neuronal activity in the delayed saccade paradigm, demonstrating a transient reset event in the SC, but not V1, leading up to saccade-related motor bursts. We then document how foveal SC neurons behave at the time of the go signal in the same task, revealing a transient bursting signal that precedes the peripheral SC reset event. Finally, we demonstrate that our observations about foveal bursts hold even in reflexive saccade tasks (without an enforced delay), resulting in the occurrence of two simultaneous short-latency neuronal activity bursts (after saccade target onset) in two disparate SC loci, one foveal and one peripheral.

### Peripheral superior colliculus neurons exhibit transient activity pauses before instructed-saccade motor bursts emerge

We analyzed peripheral SC neuron activity from a “saccades-to-gratings” task that we recently designed (8) (Materials and methods). Figure 1A, C shows the visual RF’s of two example SC neurons from this task. The saccade target (schematized by the white circle) appeared over the neurons’ RF’s, which expectedly elicited visual responses by the neurons (Fig. 1B, left and Fig. D, left); these visual responses were different for different image appearances of the saccade target, consistent with the presence of feature tuning properties in SC neurons (28, 38–40). Since the two neurons were visual-motor neurons, at the time of saccade onset later on in the same trials (Fig. 1B, right and Fig. 1D, right), both neurons also emitted a saccade-related motor burst, which could again vary in strength depending on the visual appearance of the saccade target (8, 41, 42). Remarkably, when we measured the activity of the two neurons around the time of the go signal (Fig. 1B, middle and Fig. 1D, middle), both neurons exhibited a short-latency transient pause in their activity (downward gray arrows). This pause occurred with a latency of ∼50 ms from the go signal occurrence, and it was apparently also all-or-none. That is, if the sustained activity of the neuron was high for one image feature (e.g. 0.5 cpd for the neuron of Fig. 1A, B or 11 cpd for the neuron of Fig. 1C, D), then the reduction in firing rate after the go signal was more marked than if the sustained activity was low (e.g. for 11 cpd in the neuron of Fig. 1A, B and 2 cpd in the neuron of Fig. 1C, D); this resulted in more similar firing rates after the go signal than before it, regardless of the initial firing rate that was influenced by the image appearance of the saccade target. Immediately after each pause, the strong motor burst of each neuron erupted, as classically expected from saccade-related SC neurons.

Across the population of SC neurons, and for all image features that we tested, we measured neuronal activity in the final 100 ms before the go signal. Then, we subtracted the activity of each neuron around the go signal from this baseline measurement, to obtain a baseline-subtracted firing rate (Materials and methods). When we aligned the baseline-subtracted firing rate of all neurons to the go signal, we observed a short-latency reduction in activity right before the elevation associated with saccade-related motor bursts (Fig. 2A; SC data; trials with all image features were combined together). This was consistent with what we observed from the two example neurons of Fig. 1. We also measured the raw firing rate of each neuron, both in baseline (final 100 ms before the go signal) as well as 50-150 ms after the go signal, and we compared the two measurements (Fig. 2B; trials with all image features were combined together). Once again, across the population, there was a robust reduction in firing rate after the go signal (p= 7.7693 × 10^-23^ for the spatial frequency task; p=1.4893 × 10^-19^ for the contrast task; p=6.4765 × 10^-18^ for the orientation task; signrank test).

**Figure 2.**
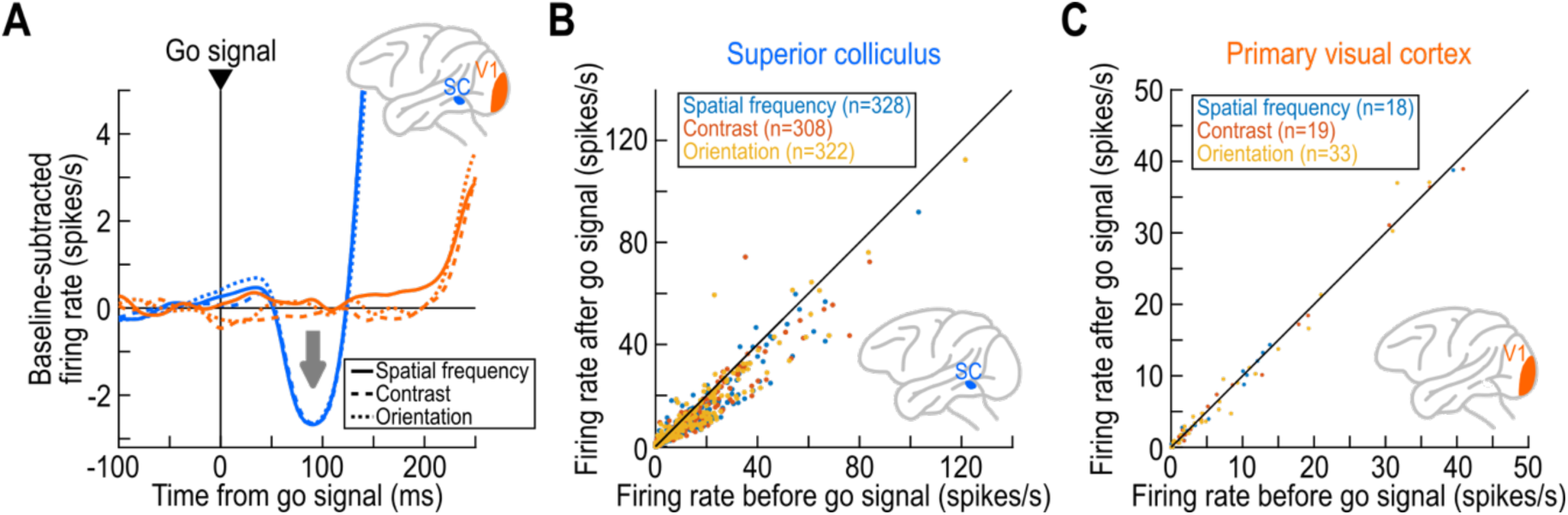
Ubiquity of SC activity pauses across task contexts, and a lack of a correlate for them in the primary visual cortex (V1) in similar behavioral tasks. **(A)** Population firing rates aligned on the go signal from the same task as in Fig. 1. Here, we baseline-subtracted the activity of each neuron based on firing rate in the final 100 ms before go signal onset (Materials and methods; numbers of neurons can be inferred from **B**, **C**). Across all three task variants (Materials and methods), SC activity pauses were present to the same extent (downward arrow), but there were no such pauses in our V1 population. The later increase in the SC curves is the saccade-related burst, and the even later V1 increase is visual reafference due to eye movement. **(B)** For each SC neuron, we measured activity in the final 100 ms before go signal onset and plotted it on the x-axis; on the y-axis; we measured average activity 50-150 ms after the go signal. For each task, there was a reduction after the go signal. **(C)** There was no such reduction in V1. Numbers of neurons are indicated in the figure, and statistical results are mentioned in the text.

Thus, we observed a short-latency pause in SC activity when releasing an instructed saccade in the classic delayed saccade paradigm (i.e. at the time of the go signal). As explained in Discusion, we think that this pause is functionally useful in the SC, especially because it represents a timely-suitable opportunity to transform SC representations from being in a visual operating mode to being in a motor one (7, 8, 10–14).

We also explored the SC pause properties further, by linking them to the sustained activity associated with the image impinging on the visual RF’s of our recorded neurons. Different image features caused different levels of sustained SC activity (28, 39) (Figs. 1B, D, 3A). If the SC pause is all-or-none, then image features with higher sustained activity should be associated with a bigger firing rate drop than image features with lower sustained activity; this would allow converging all SC firing rates to the same (low) level regardless of the image condition driving the peripheral neurons’ activity. We confirmed this by finding, for each neuron, the image condition that gave rise to the highest firing rate in the final 100 ms before the go signal, and we called this the most preferred sustained feature. We also found the image condition associated with the lowest sustained firing rate, and called it the least preferred sustained feature. Across neurons, the magnitude of the drop in firing rate between the baseline (final 100 ms before the go signal) and post-go (50-150 ms after the go signal) intervals was larger for the most preferred than least preferred sustained feature (Fig. 3B for the spatial frequency task); there was a significant difference between the two image features (p= 3.9233 × 10^-29^; t-test). Results from the contrast and orientation tasks were virtually identical, as also seen from Fig. 2A, B.

**Figure 3.**
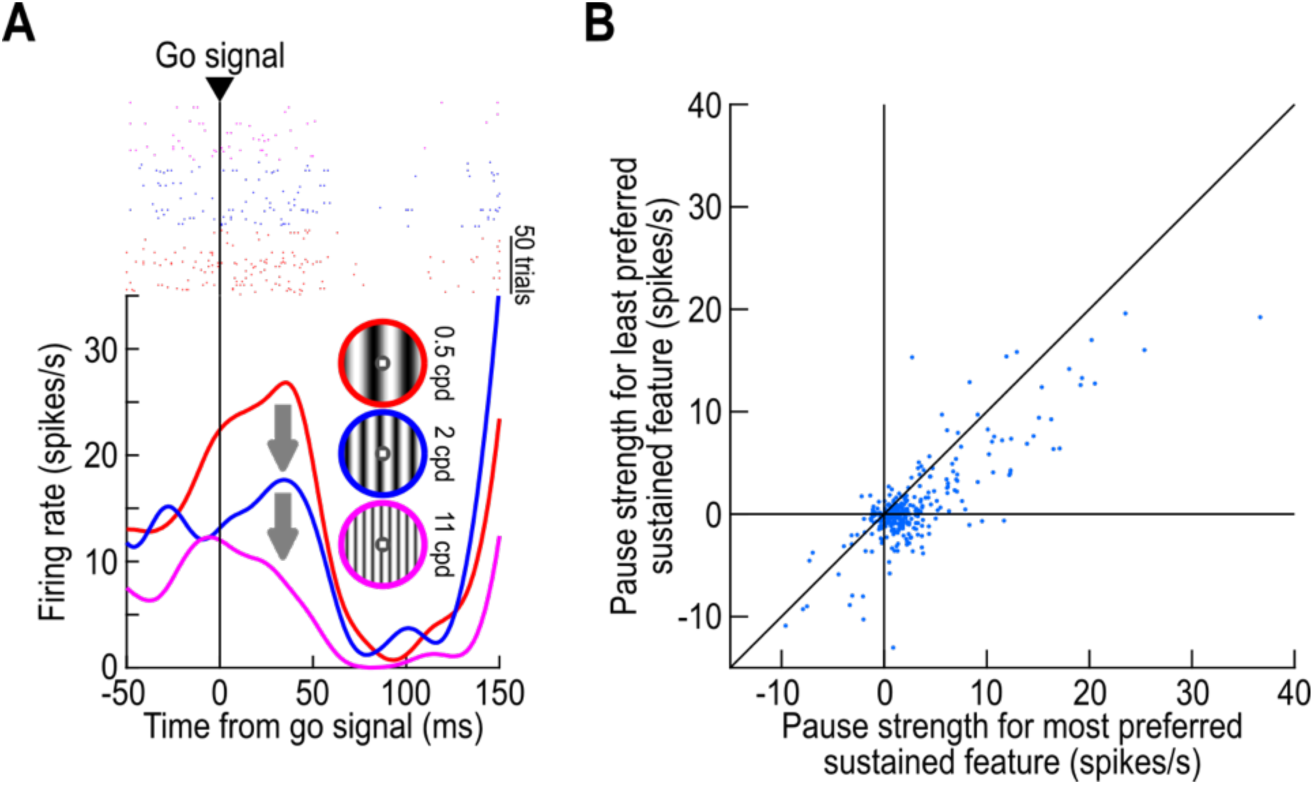
Dependence of peripheral SC activity pauses on the viewed stimulus appearance. **(A)** Activity of a third example SC neuron at the time of the go signal for three image appearances of the saccade target. During the delay period leading up to the go signal, the neuron had ongoing activity that was different for the different images, reflecting a feature tuning property of the neuron (38). As a result, the amount of activity reduction after the go signal was stimulus-dependent: the pause was stronger for the condition in which the delay-period activity of the neuron was the highest, resulting in the same low activity level regardless of initial state. **(B)** Across all neurons, we measured each neuron’s activity in the final 100 ms before the go signal, and picked the saccade-target image appearance that gave either the highest or lowest such activity. Then, we measured the pause strength (difference between activity after the go signal and activity before; Materials and methods) for the most or least preferred image feature. The pause was stronger for the most preferred image feature, consistent with **A**. Note that only data from the spatial frequency task is shown, but the other tasks revealed similar results (also seen in Fig. 2).

Thus, we identified a systematic short-latency pause in SC activity at the time of releasing a task-instructed saccade in the classic delayed saccade paradigm.

### Peripheral superior colliculus activity pauses are absent in the primary visual cortex

We also wondered whether the transient pause that we saw in the SC was a general property of visually-responsive neurons in other brain areas. Therefore, we collected a smaller number of V1 neurons from the same task (Materials and methods). There was no transient pause after the go signal, as can be seen from the V1 data in Fig. 2A. Given the fact that a large part of visual responses in the SC derives from V1 (43–48), this observation confirms that SC visual responses are functionally transformed and not merely inherited from the cortex (38, 49–52).

Also note that in the V1 neurons, there was a later elevation of neuronal activity well after the go signal (Fig. 2A), almost 100 ms later than the elevation seen for the SC neurons. This later elevation reflects visual reafferent responses in V1 as a result of eyeball rotations, whereas the earlier SC increase in firing rate (after the pause) represents the SC saccade-related motor bursts themselves. Figure 2C also shows that in V1, there was no statistically significant difference between activity right before and right after the go signal (p=0.6165 for the spatial frequency task; p=0.4209 for the contrast task; p=0.5876 for the orientation task; signrank test), suggesting an absence of a transient pause in V1 neurons.

Therefore, our results so far indicate that we observed a transient pause in SC activity at the time of the go signal in a classic delayed saccade paradigm. This pause was not a general property of other brain areas that might be recruited by the same paradigm, and that might modulate SC activity, such as V1.

### Peripheral superior colliculus activity pauses are stronger in saccade-related neurons

As mentioned above, the presence of an SC pause at the go signal might be functionally particularly useful, because it is only in the SC, and not in V1, that the very same neurons would be engaged in both visual processing and saccade-related modulations (7, 8, 10, 38, 41). If this is true, then one might expect that this pause should be more relevant for saccade-related SC neurons than for purely visual ones; this is because it is the saccade-related neurons that would exhibit both sensory and motor responses requiring a representational transformation on a rapid time scale. To check this, we classified our SC neurons as being either purely visual or motor-related. The visual neurons included visual-burst and visual-delay neurons, and the motor-related neurons included visual-motor and purely motor neurons, as defined previously (8, 38, 40).

We found that there were stronger pauses for the motor-related SC neurons than for the visual ones. For example, in Fig. 4A, B, we replotted the data of Fig. 2B but after first classifying the neurons into the two groups mentioned above. Across all tasks, there were bigger differences between the activity before and after the go signal in the motor-related neurons (Fig. 4B) than in the visual neurons (Fig. 4A). To quantify this further, we calculated a modulation index for each neuron (activity after the go signal minus before the go signal, divided by the sum; Materials and methods). This index was negative for activity pauses and zero for no pauses. In all tasks, the median value of the modulation index was more negative in the motor-related neurons than in the visual ones (Fig. 4C, D, E), and the differences between distributions of modulation indices across the two classes of neurons were statistically significant, but only marginally so for the orientation task (the results of the statistical tests are included in the legend of Fig. 4). Thus, the transient pause that we observed was stronger for motor-related neurons.

**Figure 4.**
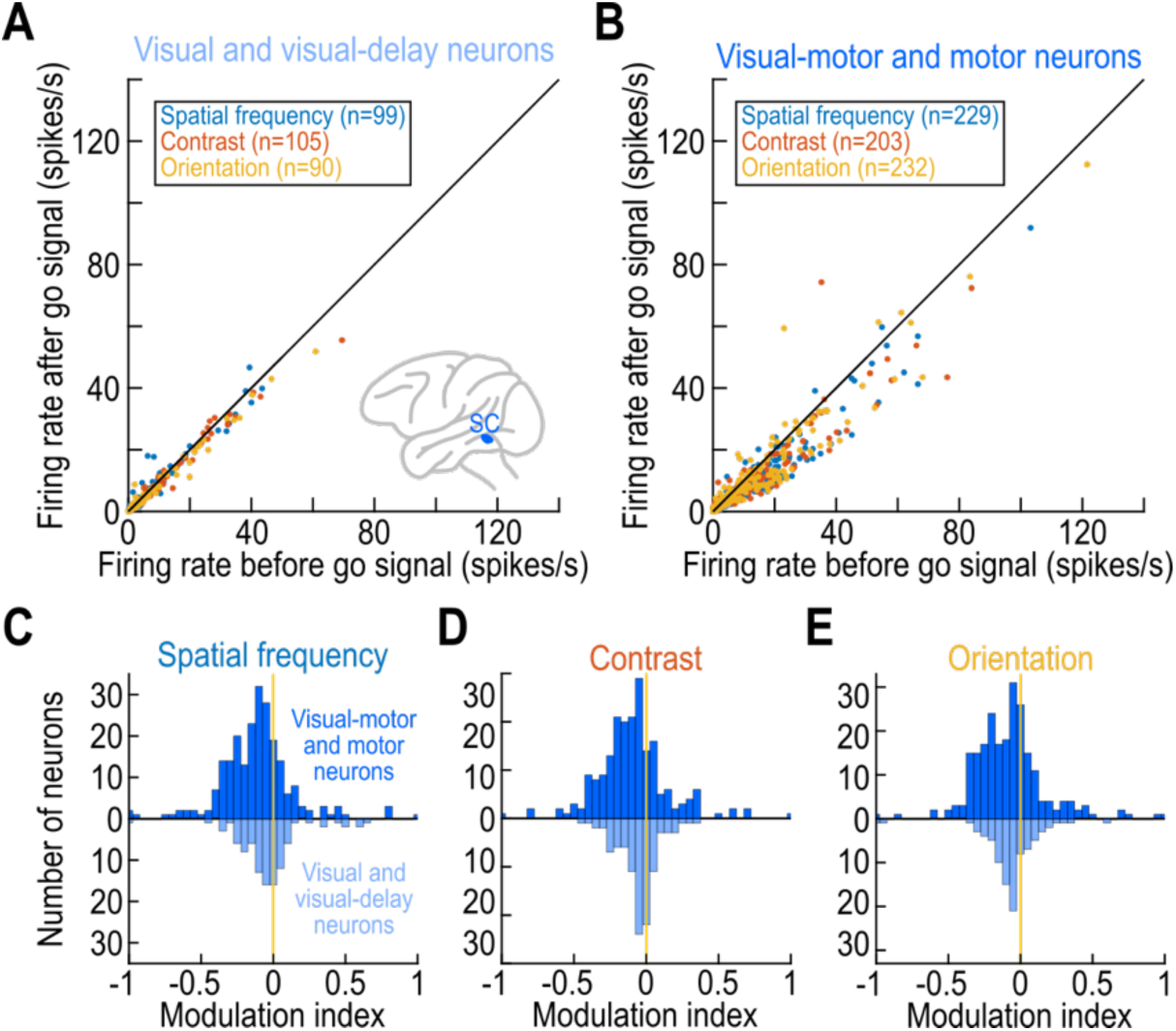
Specificity of SC activity pauses for neurons related to the motor generation of saccades. **(A)** Similar analysis to Fig. 2B, but only purely visual SC neurons. Visual neurons were visual-burst and visual-delay neurons (38). There was minimal activity reduction at the go signal in these neurons despite the presence of delay-period activity (p=9.8480 × 10^-4^ for the spatial frequency task, p=3.9107 × 10^-5^ for the contrast task, p=3.0530 × 10^-7^ for the orientation task; signrank test); also see **C**-**E**. **(B)** Neurons that needed to emit a saccade-related motor burst after the go signal underwent clearer activity reductions (p=4.5489 × 10^-21^ for the spatial frequency task, p=8.0457 × 10^-16^ for the contrast task, p=1.0135 × 10^-12^ for the orientation task; signrank test). **(C, D, E)** Neuronal modulation indices (Materials and methods) for the results in **A**, **B** across the different task contexts. Visual-motor and motor neurons had generally stronger reductions in their activity after the go signal than visual and visual-delay neurons (p=0.000532 for the spatial frequency task, p=0.004 for the contrast task, p=0.0966 for the orientation task; Wilcoxon rank sum test comparing the distribution of visual-motor/motor modulation indices to the distribution of visual/visual-delay modulation indices). The median values of the shown distributions were: -0.1069 and -0.0453 for visual-motor/motor neurons and visual/visual-delay neurons, respectively in **C**; -0.0981 and -0.036 in **D**; -0.0862 and -0.0537 in **E**.

Of course, the presence of sustained activity at the time of the go signal is a prerequisite for successfully seeing a pause, if one is present at all. Thus, it might be suggested that the differences in pause strengths that we saw in Fig. 4 between purely visual and motor-related neurons could reflect a lack of sustained activity in the visual neurons. However, this was not the case. For example, inspection of the raw firing rates on the x-axis of Fig. 4A reveals that the purely visual SC neurons had similar sustained activity levels, in general, to the visual-motor and motor neurons (x-axis in Fig. 4B). This is expected because visual neurons (especially visual-delay ones) are known to continuously represent the presence of a visual stimulus over their RF’s. Thus, the results in Fig. 4 are not explained by the purely visual neurons having no (or less) sustained activity at the time of the go signal than the motor-related neurons.

### Foveal superior colliculus neurons exhibit transient bursts instead of pauses

Having established the presence of a short-latency activity pause in SC neurons right before saccade-related bursts emerge, we next asked what happens in foveal SC representations during the delayed saccade paradigm. Figure 5A, C shows the visual RF’s of two example foveal SC neurons. In both cases, the RF’s were contained within a retinotopic eccentricity of 2 deg (dashed circle) (29), and the saccade target was at an eccentricity of 8-10 deg (Materials and methods). At stimulus onset (Fig. 5B, D, left) and saccade onset (Fig. 5B, D, right), the two neurons behaved as expected from foveal SC neurons when the saccade target is placed outside of their RF’s: at stimulus onset, there was either no modulation or a reduction in activity; and, at saccade onset, there was a strong pause when the peripheral neurons were emitting their motor bursts. Both of these observations were documented before in the SC literature (15–18), and they confirm that we were recording from foveal neurons having the saccade target location being outside their RF’s. Remarkably, right after the go signal, both foveal SC neurons showed a very strong, short-latency activity burst, which almost doubled or tripled the firing rate relative to its level at the time of the go signal (Fig. 5B, D, middle; upward gray arrows). This foveal burst occurred right before the saccade-related pause that was seen in the foveal neurons’ activity at the time of saccade onset. Thus, unlike the peripheral SC neurons (Figs. 1-4), these two example foveal SC neurons showed activity bursts, rather than pauses, when releasing instructed saccades (Fig. 5).

**Figure 5.**
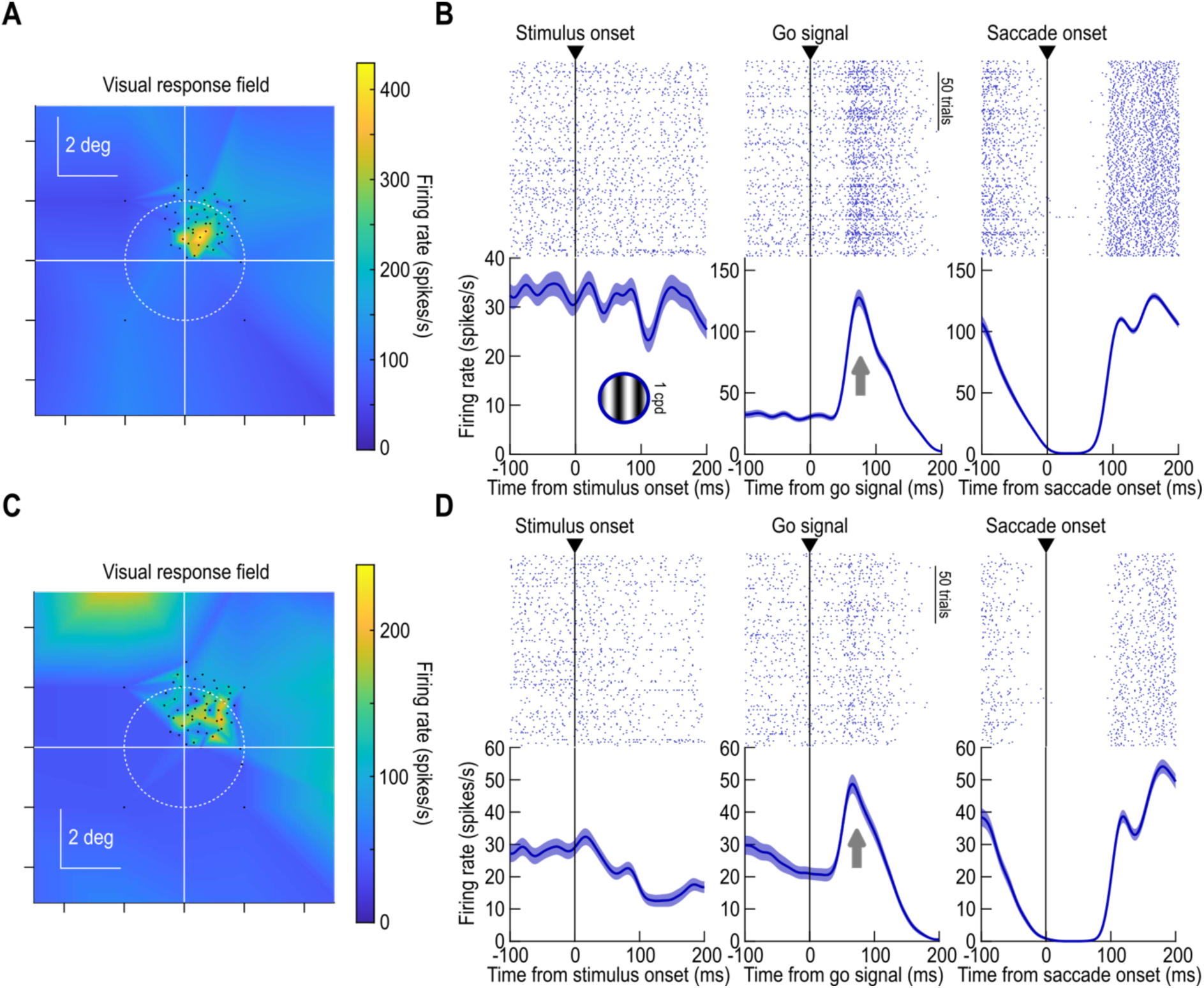
Transient activity bursts, rather than pauses, in the foveal representation of the SC prior to saccade triggering. **(A, B)** Similar to the formatting of Fig. 1 but now for an example foveal SC neuron. The RF was constrained in the central 2 deg of visual angle (**A**), and the saccade target was at a farther eccentricity (15). In **B**, the neuron behaved as expected at the times of stimulus (left) and saccade (right) onset; these observations confirm that the saccade target (and target vector in the case of the motor epoch) was outside of the neuron’s RF. After the go signal (middle), the neuron showed a strong transient burst in activity, and at a qualitatively similar time to the peripheral pauses of Figs. 1-4. Note that this neuron is the same example neuron of Fig. 1 in (15), but here we additionally showed the activity at the time of the go signal. **(C, D)** Similar observations for a second example foveal SC neuron. There was again a transient burst right after the go signal, in addition to expected classic stimulus- and saccade-related modulations. All other conventions are like Fig. 1. Error bars denote SEM across trial repetitions. Panel **A** was adapted from (15).

Just like with the peripheral neurons, we also had multiple visual features of the eccentric saccade target in the version of our delayed saccade paradigm that we used when recording our foveal SC neurons (Materials and methods). This allowed us to investigate whether there was any stimulus dependence of the foveal activity bursts. In Fig. 6A, the foveal burst of a third example foveal SC neuron is shown, but this time for the two visual appearances of the eccentric saccade target. In both cases, the strength of the foveal burst was the same. This was a general property across our population of foveal SC neurons. Specifically, we measured average firing rate 50-100 ms from go signal occurrence, and we did so for either the low or high spatial frequency saccade target. There was no difference in foveal burst strength (p=0.8509; signrank test across the population; n=33). Therefore, unlike in the peripheral neurons, the foveal SC burst was not stimulus-dependent. As we will show later with our timing analyses, this burst might thus be a general trigger signal to aid in jumpstarting peripheral saccadic orienting, regardless of the peripheral target’s image appearance.

**Figure 6.**
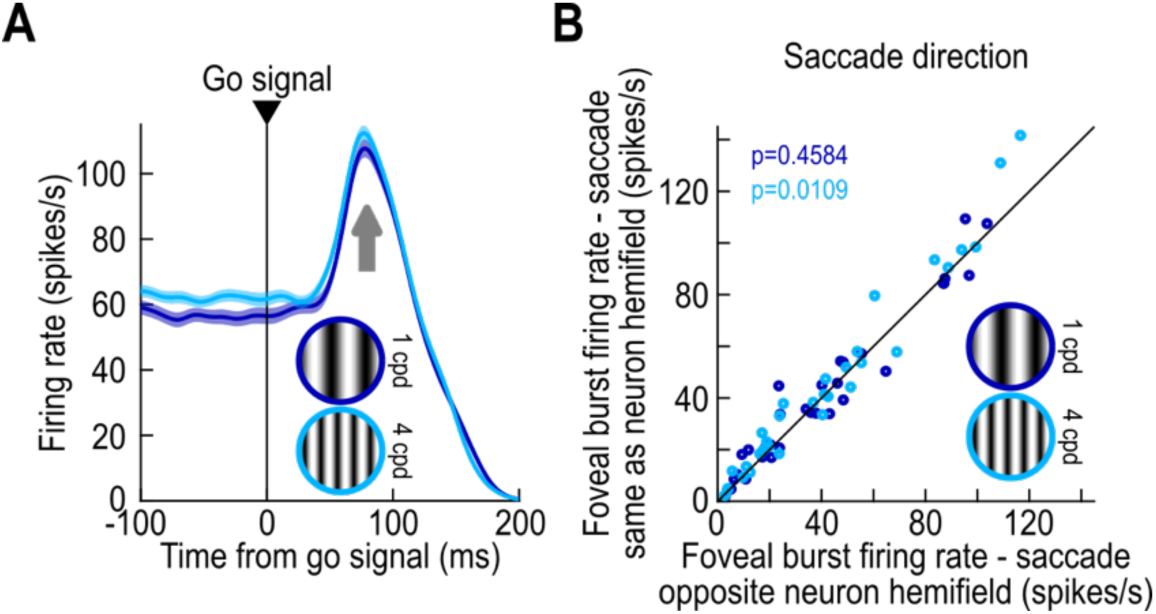
Lack of stimulus or direction dependence of the foveal bursts. **(A)** The foveal burst of an example SC neuron for two visual appearances of the peripheral saccade target. The foveal burst was similar in strength for the two stimuli. Error bars denote SEM across trials. **(B)** We also checked whether the foveal burst depended on congruence between the recorded foveal SC hemifield and the saccade direction. There were no or moderate dependencies on saccade direction (signrank test; n=33).

We also checked whether the foveal SC burst depended on saccade direction. Specifically, foveal SC neurons are lateralized (29), just like peripheral ones are. Thus, if a foveal neuron was recorded from the right SC, then it represented the left foveal space. In this case, a leftward eccentric saccade target would elicit motor bursts in the peripheral neurons of the same SC as the recorded foveal neuron, whereas a rightward saccade would elicit motor bursts in the opposite SC. As can be seen from Fig. 6B, there was no clear dependence of foveal burst strength on saccade direction; for the high spatial frequency saccade target, there was a statistically significant result, but the magnitude of the difference between hemifield directions was not qualitatively different from that seen with the low spatial frequency target (for which there was no statistically significant difference between saccade directions). Thus, foveal SC bursts (Figs. 5, 6A) also did not show a systematic dependence on saccade direction relative to the recorded neurons’ RF hemifields (Fig. 6B).

### Foveal superior colliculus activity bursts lead peripheral pauses

Our results so far indicate that releasing an instructed saccade in the classic delayed saccade paradigm is associated with a short-latency (foveal) burst in one part of the SC topographic map and a (more peripheral) pause in another (Fig. 7A). To understand the potential links between these two types of neuronal responses further, we summarized SC population activity in the two groups of neurons, after normalizing the activity of each neuron to its activity in the final 100 ms before go signal onset. At the time of saccade onset (Fig. 7B), the peripheral neurons emitted strong saccade-related motor bursts, and the foveal neurons decreased their activity, consistent with the example neurons of Fig. 5 and with the prior literature (15–17). However, this relationship was completely reversed in the go signal epoch (Fig. 7C), with foveal neurons now exhibiting an approximately three-fold increase in their activity immediately after the go signal and right before saccade triggering; this is consistent with the foveal burst strengths seen in the example neurons of Fig. 5. The peripheral neurons, on the other hand, paused. Interestingly, plotting the peripheral and foveal neurons together in Fig. 7C revealed that the foveal burst actually led the peripheral pause by some time. To better quantify this timing relationship, we binned spike times in all neurons into 2 ms non-overlapping time bins, and we estimated spike likelihood in each time bin after go signal onset. The onset of the foveal burst (defined as the first time point to have at least three successive increases in spike likelihood; Materials and methods) led the onset of the peripheral pause (defined as the first time point to have at least three successive decreases in spike likelihood; Materials and methods) by 10 ms.

**Figure 7.**
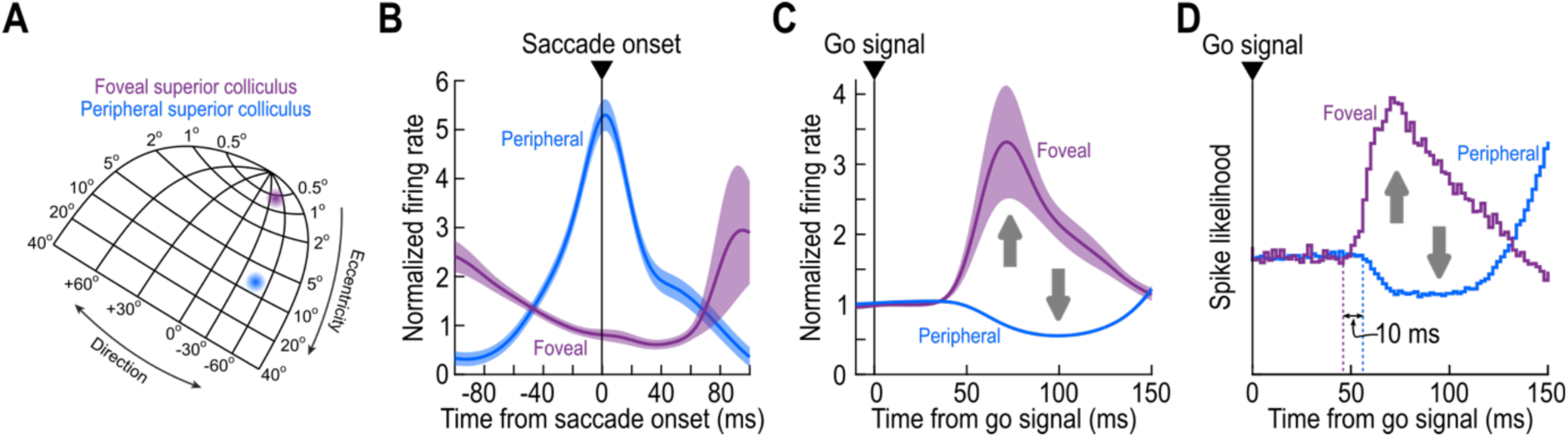
Temporal sequencing of foveal bursts and activity pauses in the SC prior to saccade triggering. **(A)** Schematic of the SC topographic map (29, 53, 54), with an example indication of the relative positions of neurons that were recruited in our populations of recordings from either the experiments of Figs. 1-4 or those of Figs. 5-6. **(B)** At the time of saccade onset, peripheral neurons exhibited a motor burst, and foveal neurons reduced their activity, as expected (6, 15–17, 30, 55). Error bars denote SEM across neurons, and the normalization factor for each neuron was the activity level in the final 100 ms before go signal onset; see **C**. **(C)** At the time of the go signal, foveal neurons exhibited strong bursts, which started slightly earlier than the peripheral SC pauses. **(D)** We quantified this timing differences by having 2 ms bins in which we measured spike likelihood across all neurons. Then, we identified the onset of the burst or the pause as the first time point at which the spike likelihood changed in the same direction (increase for bursts and decrease for pauses) for at least three more upcoming time bins (Materials and methods). The peripheral pauses lagged the foveal bursts by 10 ms.

Thus, foveal SC bursts at the time of releasing instructed saccades in the classic delayed saccade paradigm occur systematically earlier than peripheral SC pauses.

### Foveal superior activity bursts reflect instructed saccade context

A parsimonious explanation for our results so far could relate to intrinsic RF properties of SC neurons (38), combined with potential long-range lateral inhibition mechanisms (19–24). Specifically, it could be possible that the removal of the fixation spot in the delayed saccade paradigm may act as an offset sensory stimulus to the foveal SC neurons (i.e. the fixation spot in this case is a visual stimulus that is positioned somewhere within the foveal RF’s, and its offset is a sensory event that could lead to a sensory offset response). Since SC neurons are expected to sometimes have sensory offset responses (i.e. activity bursts in response to an offset of a visual stimulus that was previously present in their RF’s) (38, 56–59), it is conceivable that the fixation spot removal in our saccade paradigm could trigger an offset response in foveal SC neurons. After such a response, lateral inhibition in the SC might, in turn, cause peripheral neuron activity pauses (as a result of the foveal bursts). To investigate this potential cascade of events, we first compared our foveal bursts to real offset responses as measured by our RF mapping tasks. We then explored other situations with offsets of visual stimuli in our paradigms, but without an explicit task instruction to release a saccade. And, finally, we explored cases in which we created a competition between putative lateral inhibition and the occurrence of peripheral SC activity bursts (rather than pauses), to understand whether the peripheral pauses are due to lateral inhibition from the foveal bursts or not. We describe the results of these three successive investigations next.

To explore whether foveal bursts at the go signal reflect offset responses to the removal of the fixation spot, we measured offset responses in our foveal SC neurons from our RF mapping tasks, which also used similar small white spots as the stimuli (Materials and methods). During RF mapping, we presented a small white spot (similar to the fixation spot) at different locations near the fixation spot that the monkeys were looking at. After a few hundred milliseconds, the white spot was removed, allowing us to measure sensory offset responses. Figure 8A, B shows the offset responses of the two example foveal neurons of Fig. 5. Because foveal SC RF’s are strongly lateralized (29) (Fig. 5A, C), fixation spot removal in our main task was equivalent to an offset response for a stimulus that was visible at a non-zero distance, *r*, from the RF hotspot eccentricity (assuming, rightly, that the monkeys properly centered gaze on the fixation spot, on average); here, *r* would be the distance between the RF hotspot and the central preferred retinal locus of fixation. Therefore, from the RF mapping data, we picked a ring of sampled stimulus locations at a distance *r* from the RF hotspot (+/- 0.25 deg; Materials and methods). We took all of these stimulus locations in the hemifield of the RF hotspot (to maximize the likelihood of seeing offset responses), and we plotted the RF offset responses in Fig. 8A, B. As can be seen, at the same eccentricity from the RF hotspot as the foveal fixation spot in the main task, the two example neurons did not emit substantial offset responses at all. For comparison, the insets in Fig. 8A, B replicate the foveal bursts of Fig. 5 from the same neurons, showing that the neuronal responses at the go signal in the main task were very different from their responses for the offset of small white spots near their RF hotspots. Thus, for these two example neurons, foveal bursts were not trivially explained by sensory offset responses to the removal of the foveal fixation spot.

**Figure 8.**
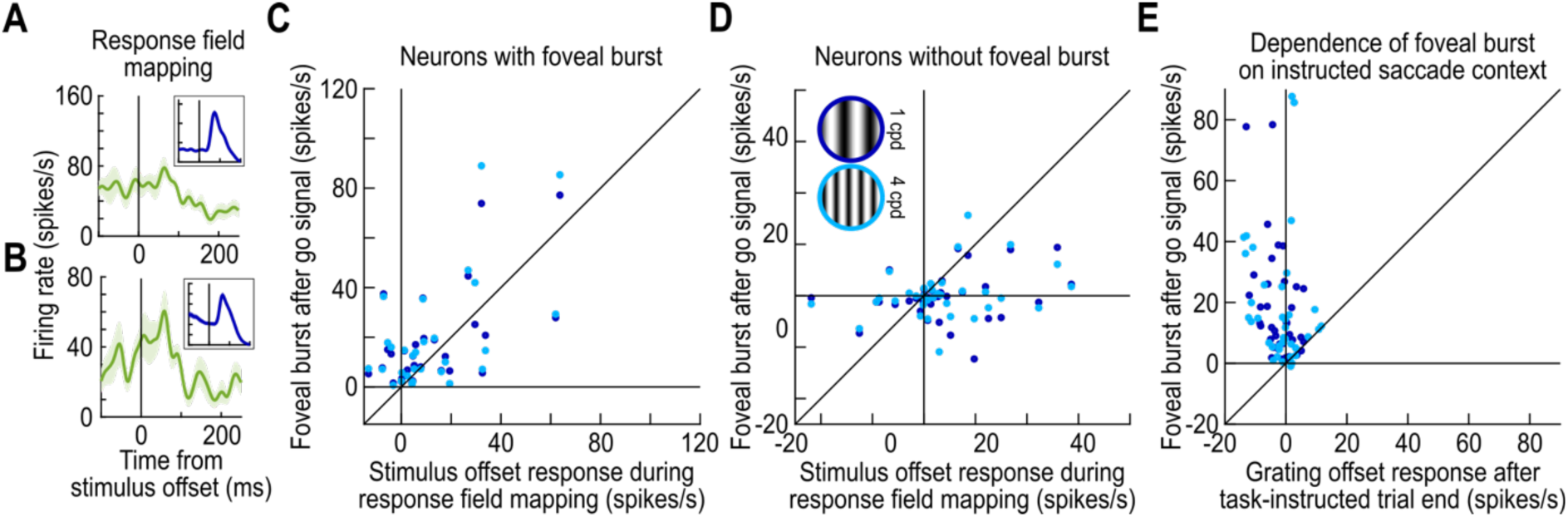
Dissociation between foveal bursts after the go signal and stimulus-offset responses in the foveal SC neurons. **(A)** For the example neuron of Fig. 5A, B, we measured the response of the neuron to the offset of a small spot during the RF mapping task (Materials and methods); we took care to define a band of eccentricities and directions in which the stimulus location was in a similar portion of the RF as the disappearing fixation spot in the main task (Materials and methods). The neuron did not exhibit a substantial offset response; for comparison, the inset shows the foveal burst of the same neuron from Fig. 5B. Thus, the foveal burst was not a sensory offset response to the disappearance of the fixation spot in our saccade task. Error bars denote SEM across trial repetitions. **(B)** Similar observations for the second example neuron of Fig. 5C, D. **(C)** For each neuron that exhibited a foveal burst in the main saccade task (Materials and methods), we plotted the foveal burst strength against the sensory offset response of the neuron from the RF mapping task (Materials and methods). There was a moderate relationship between sensory offset responses and foveal bursts (r = 0.6026, p = 0.0005 for the low spatial frequency; r = 0.5924, p = 0.0007 for the high spatial frequency). Each color shows the foveal burst for a one spatial frequency of the saccade target. **(D)** For neurons with no foveal burst in the main saccade task (Materials and methods), they could still exhibit offset responses in the RF mapping task (r = 0.2662, p = 0.1166 for the low spatial frequency; r = 0.2890, p = 0.0874 for the high spatial frequency). Thus, **C** and **D** combined suggest that foveal bursts were not always explained by sensory offset responses due to the disappearance of the fixation spot in the delayed saccade paradigm. **(E)** In the main saccade task, after a successful saccade to the grating, the grating (which was now foveal) eventually disappeared from the foveal RF’s, and the monkey was free to make any saccades in the inter-trial interval. The foveal neurons did not burst for the disappearance of the grating, but they still burst for the disappearance of the fixation spot during the instructed-saccade phase of the trial (p = 6.6424 × 10^-8^ for the low spatial frequency, p = 1.5453 × 10^-7^ for the high spatial frequency; signrank test). Thus, foveal bursts were task-dependent.

Across the population of foveal SC neurons, we then collected sensory offset responses and compared them to foveal bursts. In Fig. 8C, we took the neurons that statistically exhibited foveal bursts at the go signal (Materials and methods), and we investigated how they behaved in terms of sensory offset responses during the RF mapping task. The neurons did generally exhibit offset responses, as might be expected (38). Importantly, we also looked at foveal SC neurons that did not statistically emit foveal bursts in the main delayed saccade task (Materials and methods). If our foveal bursts of Figs. 5-7 were fully explained by RF offset responses, then these neurons should not have exhibited any offset responses at all in the RF mapping task. In reality, this was not the case at all (Fig. 8D). There were clear offset responses in the RF mapping task for the neurons that did not exhibit strong foveal bursts at the go signal in the main delayed saccade task. Thus, foveal bursts at the go signal in the classic delayed saccade paradigm could be distinct from simple sensory offset responses associated with fixation spot removal.

Perhaps the strongest evidence that we had for a dissociation between offset responses and the foveal SC bursts at the go signal came from our main delayed saccade task itself. Specifically, in this task, after the monkeys foveated the instructed saccade target (the spatial frequency grating), this target eventually disappeared after a few hundred milliseconds of fixation (Materials and methods). Since the monkeys had successfully foveated the saccade target prior to that, the removal of this target in this case was equivalent to an offset stimulus for the neurons’ RF’s. Critically, the monkeys had no explicit task instruction associated with the target removal; this disappearance of the visual stimulus was the onset of the inter-trial interval in which the monkeys were free to look wherever they wanted (Materials and methods), and we analyzed neuronal activity of the foveal neurons upon target removal only when there were no saccades for at least 150 ms after the removal. As can be seen from Fig. 8E, the neurons that did show foveal bursts at the go signal (y-axis) did not show substantial activity elevations upon (foveal) target removal (x-axis).

Thus, all of the analyses of Fig. 8 suggest that foveal SC bursts are dependent on the task context, and may not trivially reflect simple stimulus-offset responses due to fixation spot removal in the classic delayed saccade paradigm.

### Foveal bursts cooccur with peripheral visual responses in reflexive saccade tasks

If foveal bursts are indeed not explained by stimulus offset responses, then are the peripheral SC pauses at least still explained via long-range lateral inhibition mechanisms in the SC (19–24)? To test for this, we employed another classic saccade-related task, now not enforcing a delay period between fixation spot removal and eccentric saccade target appearance (Fig. 9A). When recording from peripheral SC neurons in this task, short-latency visual responses appear, before the saccade-related motor bursts quickly evolve (50, 60–63). If lateral inhibition was the sole determinant of complementary transient pauses and bursts at disparate SC loci (Fig. 7), then the peripheral visual bursts in this version of the saccade task might be expected to eliminate the foveal bursts that we observed above, and maybe even replace them with activity pauses instead. Thus, we recorded not only peripheral SC neurons, but also foveal ones in the classic reflexive, visually-guided saccade task (Fig. 9B).

**Figure 9.**
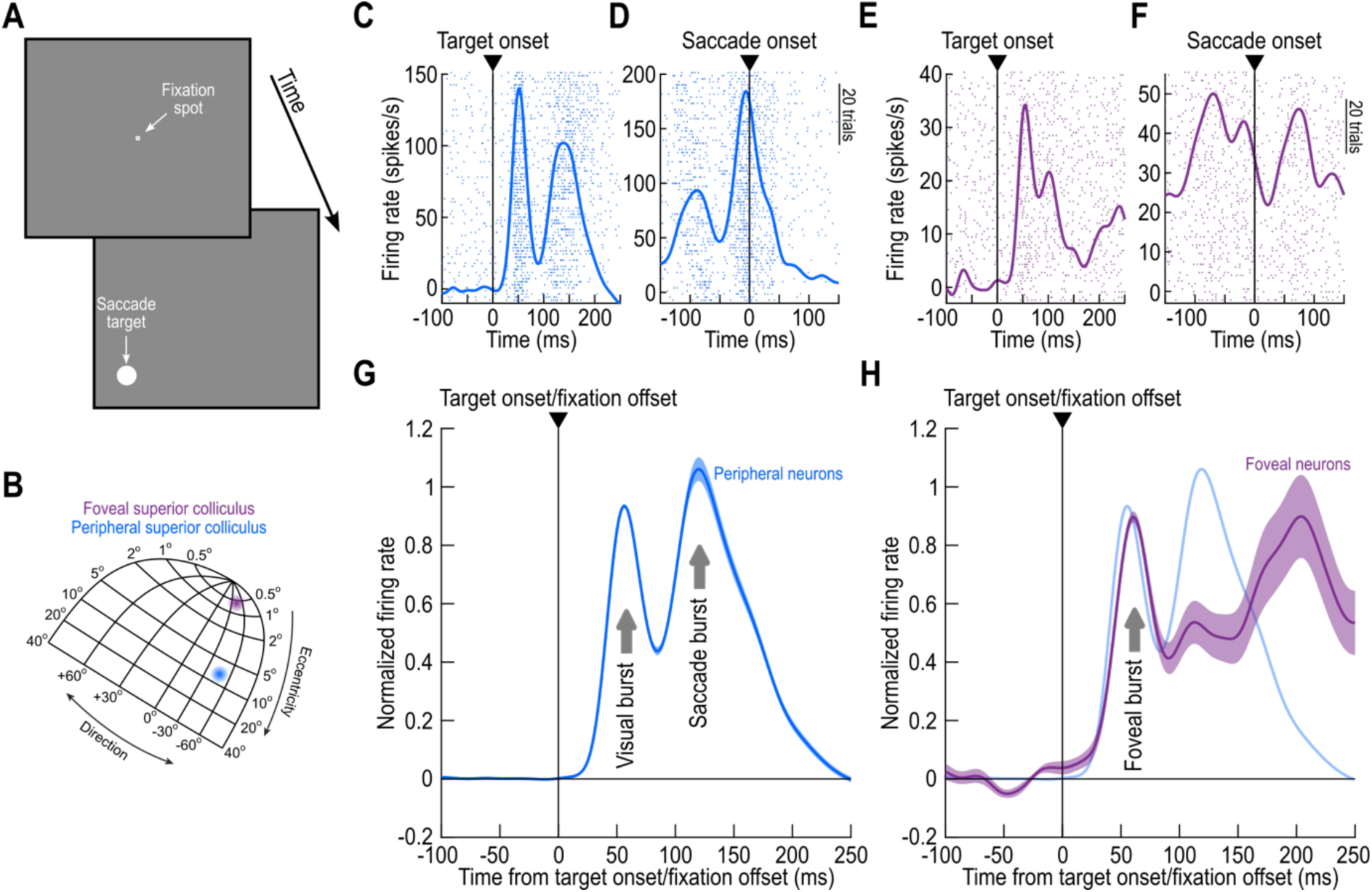
Persistence of foveal bursts even in immediate, visually-guided saccade tasks. **(A)** In this task, the go signal (fixation spot disappearance) coincided with the onset of the peripheral saccade target. **(B)** Thus, we could ask whether at the time of the peripheral visual burst in response to target appearance, we could still observe a foveal burst. **(C, D)** Example peripheral SC neuron showing expected activity discharge: there was an initial visual burst (**C**), followed immediately by a saccade-related motor burst (**D**). **(E, F)** Example foveal SC neuron from the same task. Remarkably, the foveal neuron burst (**E**) at the same time as the peripheral visual burst in **C**. It then reduced its activity at saccade onset (**F**), as expected from foveal SC neurons when eye movements are triggered (15, 17, 55). Trial numbers in **C**-**F** can be inferred from the spike rasters, and error bars denote SEM across trials. **(G)** Population results for all peripheral neurons collected during this task. Error bars denote SEM across neurons. Visual and motor bursts were evident. **(H)** For the foveal neurons, they exhibited a foveal burst even when the peripheral neurons were bursting for the visual onset inside their RF’s (the faint blue curve is a replication of the curve in **G** to illustrate the similarity of the timing of the two bursts). The neurons then reduced their activity when the peripheral neurons were bursting at saccade onset (post-saccadic reafferent responses also emerged). Thus, in the immediate, visually-guided saccade task, there are two simultaneous bursts at two different loci in the SC after target appearance. Error bars denote SEM across neurons.

Figure 9C shows the activity of an example peripheral SC neuron. Here, we plotted the neuron’s activity aligned on eccentric saccade target onset (which coincided with fixation spot removal in this task; Fig. 9A), and after subtracting pre-stimulus activity like we did recently (25). The neuron exhibited a short-latency visual response, followed <50-100 ms later by a second volley of spiking. This second volley of spiking was the saccade-related motor burst, as can be seen when aligning the same data to saccade onset (Fig. 9D). Our prior work with this task has taught us that the same rapid transformation from a visual to a motor regime also does take place in it (8), suggesting that a foveal burst signal might still be recruited in such a task. Remarkably, this was indeed the case. When we recorded from an example foveal SC neuron in this task, it exhibited a strong foveal burst (Fig. 9E), before it expectedly paused at saccade onset (Fig. 9F). The net result was that the two example neurons of Fig. 9C-F provided suggestive evidence for the existence of two simultaneous short-latency activity bursts in the SC at two very disparate loci: one peripheral representing the saccade target appearance, and one foveal associated with the go signal for generating a saccade.

Across the population, our results were consistent with those seen from the two example neurons. Peripheral SC neurons showed expected biphasic responses consisting of a first visual burst followed by a later saccade-related motor burst (Fig. 9G). And, remarkably, foveal SC neurons showed a transient burst at the same time as the peripheral visual bursts (Fig. 9H), before reducing their activity at the time of saccade onset. Note that in this task, these foveal bursts occurred for non-grating stimuli (Materials and methods), indicating that our earlier foveal burst results above were not a property of the visual appearance of the eccentric saccade target. Also note that these foveal bursts were not artifacts of us erroneously recording the inner edges of peripheral RF’s. If that was the case, we should have seen saccade-related bursts in these neurons (albeit weak), but we did not; we also checked the RF’s of our neurons (Materials and methods).

Thus, the complementary nature of transient pauses and bursts at two disparate SC loci that we saw in the delayed saccade paradigm above (e.g. Fig. 7C) was not an obligatory observation (as would be dictated by lateral inhibition mechanisms). Clearly, simultaneous short-latency bursts are possible (Fig. 9). Interestingly, we also previously saw that it is possible to observe simultaneous activity increases in two different locations on the SC map (64) Similarly, prior experiments with peripheral SC transient pauses concluded that they were not fully explained by lateral inhibition from foveal SC responses to foveal image changes (65).

### Peripheral visual responses, but not foveal bursts, correlate with saccadic reaction time

The immediate saccade paradigm of Fig. 9A also allowed us to further explore additional properties of foveal SC bursts. Specifically, we asked to what extent these bursts might relate to saccade timing variability. In peripheral neurons, we replicated the expected finding that SC visual responses are significantly stronger (and earlier) for faster saccadic reaction times (28, 50, 51, 60–63). This can be seen from the population results shown in Fig. 10A, C: here, we split the trials for each neuron according to whether the saccadic reaction time was faster or slower than the median of the session, and we found that the faster trials had stronger peripheral SC visual bursts (Fig. 10A, C). No such relationship emerged for the simultaneously occurring foveal SC bursts (Fig. 10B, D), again suggesting that these bursts might be a switch-like signal that is independent of saccade-target appearance (Fig. 6A), saccade direction (Fig. 6B), or saccade timing (Fig. 10B, D). The results of Fig. 10B, D also confirm that these transient responses in the foveal SC were not responses from the inner edges of eccentric SC RF’s associated with the target location; if that was the case, we should have seen results like in Fig. 10A, C, as well as saccade-related bursts at the time of saccade onset.

**Figure 10.**
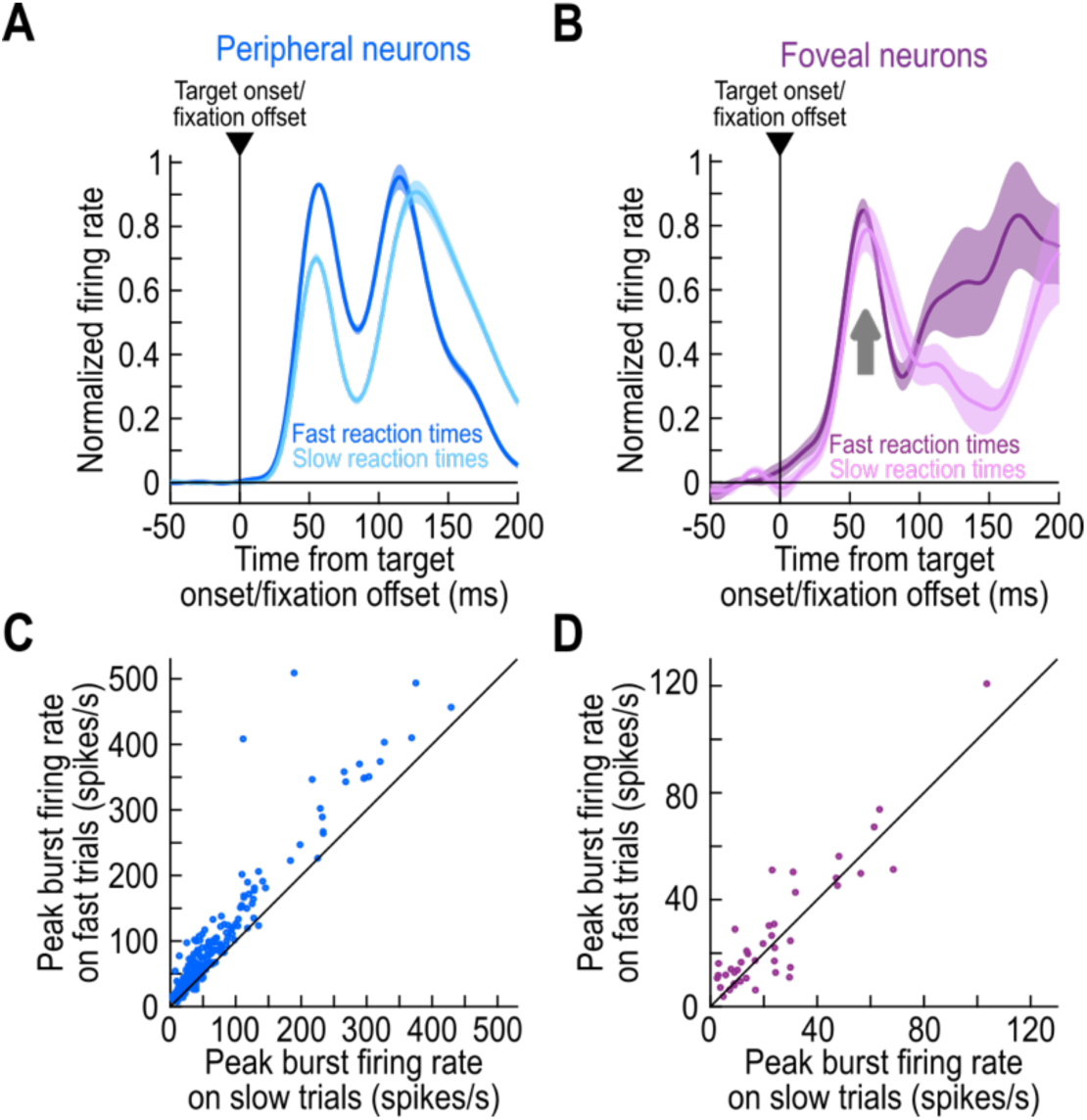
Dissociation between foveal bursts and saccadic reaction times in immediate, visually-guided saccade tasks. **(A)** In peripheral SC neurons, we replicated expectations from the previous literature (28, 50, 51, 60–63): visual bursts in response to target onset were weaker and slightly later for later saccadic reaction times. In this figure, we split trials by the median saccadic reaction time of each session. Then, we plotted the population firing rates. Trials with late saccades (light blue; later saccade bursts than in the saturated blue) had much weakened and slightly delayed visual bursts. Error bars denote SEM across neurons. **(B)** In the foveal neurons, we did not observe a clear difference in foveal bursts between trials with fast and slow saccadic reaction times (the different times of post-burst activity reductions in the two curves reflect the different times of saccade onsets in the two sets of trials per neuron). Error bars denote SEM. **(C)** Individual neuron results from the analysis of **A**. Visual bursts were stronger on fast trials (p = 2.6741 × 10^-35^; signrank test). n=253 neurons. **(D)** For foveal neurons, there was no difference (p=0.059; signrank test). n=41 neurons.

Finally, and for completeness, we also revisited the foveal bursts from our delayed saccade paradigm. When we analyzed these bursts as a function of saccadic reaction time, we again found that the bursts had the same strength for either the fast or slow saccadic reaction time trials (Fig. 11). Peripheral SC pauses also did not appear to have a systematic relationship to saccade timing in the same task, which might be expected given the task structure of prior motor preparation associated with the delay period.

**Figure 11.**
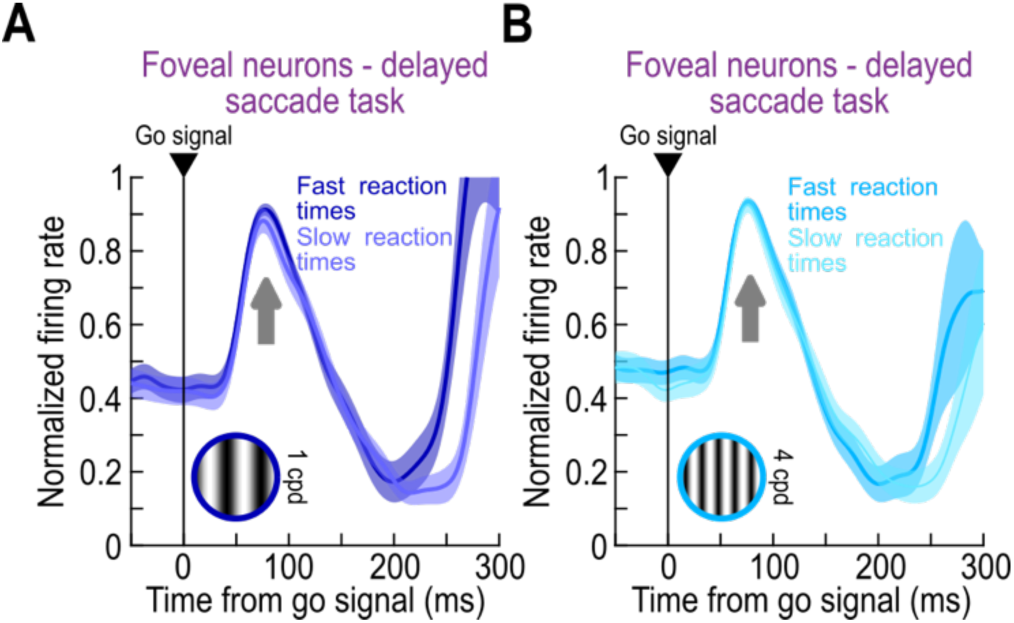
Dissociation between foveal bursts and saccadic reaction times also in delayed saccade contexts. **(A)** For the delayed saccade task of Figs. 5-8, we split trials based on the median saccadic reaction time of a given session (just like we did for Fig. 10), and we then plotted population firing rates for the two split groups. Foveal bursts were similar in both cases. **(B)** Similar results for the other spatial frequency of the peripheral saccade target. Thus, the foveal bursts behaved similarly in both versions of our visually-guided saccade paradigm in terms of subsequent saccade timing. Error bars denote SEM across neurons.

## Discussion

We investigated neuronal response dynamics at the time of “releasing” instructed saccades in two classic visually-guided saccade paradigms. Such paradigms are routinely used in the study of primate sensation, cognition, and action. We specifically identified a transient foveal SC signal that precedes the actual act of peripheral saccadic orienting by a short time. In delayed saccade paradigms, this foveal signal, which is time-locked to the go signal of the paradigms, leads peripheral SC activity pauses (again time-locked to the same go signal) by ∼10 ms. This foveal signal is also not explained by stimulus-offset responses, does not depend on stimulus appearance or saccade direction, and is independent of eventual saccade timing. Remarkably, in immediate saccade paradigms, this transient foveal signal still occurs, resulting in two simultaneous activity bursts at two disparate loci in the SC topographic map.

Peripheral SC activity pauses have been reported before. For example, Li and colleagues employed a foveal stimulus change as a cue in a target selection paradigm. Peripheral SC neurons exhibited transient pauses in their activity at the time of the foveal stimulus change (65) Similar observations were made in a variety of related tasks, most of which involving the appearance or modification of a sizeable foveal visual stimulus, and in multiple brain areas like the SC, frontal eye fields (FEF), and lateral intraparietal area (LIP) (37, 66–71); in some of these studies, the removal of the fixation spot was accompanied by the onset of multiple peripheral stimuli at the same time, which could contribute to the pauses (72). Here, we explicitly recorded foveal SC neurons without presenting a new foveal stimulus and demonstrated that peripheral SC pauses might be related to a transient foveal signal. Importantly, this signal is likely not a simple sensory response, because it clearly depended on the task context, and also because it was dissociated from stimulus-offset responses observed during RF mapping (Fig. 8). Moreover, the foveal signal need not directly mediate peripheral pauses via lateral inhibition (19–24). This is because we saw peripheral visual bursts at the same time as the foveal bursts in the immediate saccade paradigm (Figs. 9, 10); in this case, it would be difficult for lateral inhibition mechanisms alone to account for such simultaneity of bursts.

We believe that there are at least two good reasons for transient peripheral SC pauses to occur in our investigated paradigms. The first is biophysical. SC saccade-related bursts are explosive, and can exceed 1000 spikes/s (73). Thus, a transient pause right before motor bursts could aid in the biophysical processes leading up to burst generation (74–76).

The second reason why peripheral SC pauses may be functionally useful is that a large number of saccade-related SC neurons are also sensory neurons as well (31, 38). This requires a representational transformation between visual and motor regimes (7–10, 77), and this transformation even needs to happen within only a few tens of milliseconds in the immediate saccade paradigm (8). For example, Jagadisan and Gandhi (7) showed that while visual and motor bursts in the SC can appear qualitatively similar to each other and reach similar firing rates, the temporal structure of SC population activity is altered at the time of saccade generation. In our subsequent confirmation of this observation, we also noted that activity subspaces could be orthogonal to each other in the two neuronal processing regimes, and that individual neuron preferences for specific images can change between visual and motor epochs (8). Thus, peripheral SC pauses right before the motor bursts would constitute a perfect resetting mechanism for rapid representational transformations to be implemented. It would be interesting in future studies to understand how these transformations themselves emerge.

In terms of the fovea, prior work has recorded from the deep rostral SC during saccades. Consistent pauses in foveal SC activity were observed at the time of saccade generation (17, 18). Interestingly, prior to saccade generation, there were still some hints in some of these studies for subtle elevations in foveal SC activity when releasing saccades by certain task events (16, 19, 32–37). However, there was nothing reported that was as strong as we saw, and the foveal elevations were not the focus of these earlier studies (and thus not exhaustively characterized).

From the perspective of the peripheral reset alluded to above, our observed foveal bursts are particularly interesting. This is because they may act as the trigger signal jumpstarting peripheral pausing, and thus actively participate in the orienting process. Consistent with this, our foveal bursts were not stimulus or saccade-direction dependent. They were also not related to subsequent saccade timing, unlike (peripheral) visual bursts (50, 60–63). And, most importantly, they still occurred in the immediate saccade paradigm, which still requires a representational transformation between visual and motor regimes (8). Of course, the question is: what drives these bursts? In our work, we tried to dissociate them from simple offset responses due to the removal of the fixation spot. Interestingly, we found that target removal at the ends of trials in the main delayed saccade paradigm was not associated with foveal bursts, suggesting that the bursts were indeed dependent on the context of subsequently releasing a planned, instructed saccade.

More broadly, we think that it is not unheard of for a foveal signal in the SC to be relevant for peripheral SC processes. For example, we recently found, in the context of peri-microsaccadic changes in peripheral visual sensitivity, that exclusive experimental control over foveal SC state is sufficient to modulate peripheral visual sensitivity (25). Similarly, in the opposite direction, we also found that foveal SC state can be modulated by peripheral SC state across saccadic eye movements (15). These observations may relate to wider concepts like traveling waves. While such waves are presently gaining more research interest, classic SC work has indeed demonstrated a potential role of such waves in active vision (78, 79). In the future, it would be interesting to investigate such waves in more detail. For example, we can attempt to understand the links between foveal bursts and peripheral pauses by blocking the Inputs to the SC from the cortex and investigating whether signatures of an impact of foveal bursts still appears in the peripheral SC representation after losing cortical drive.

We are also especially intrigued by our observation of simultaneous bursts in the foveal and peripheral SC in the immediate visually-guided saccade paradigm. These simultaneous bursts are significant because they suggest that the peripheral pauses in the delayed saccade paradigm may not necessarily result from lateral inhibition mechanisms. In fact, in the immediate saccade paradigm, it was known for many decades that peripheral visual bursts are actually enhanced, rather than suppressed, relative to visual bursts during fixation (80). Since it is very likely that foveal bursts still occurred in these classic experiments (80), had they been investigated, it remains to be seen whether the foveal bursts may actually aid in the peripheral enhancement. Certainly, in our recent work, we found that enhancing pre-stimulus activity in the foveal SC can indeed help in enhancing peripheral visual bursts (25). We should also note that there are other observations in the literature for which direct evidence of lateral inhibition between the foveal and peripheral SC representations was absent. For example, when we investigated microsaccade generation in the foveal SC representation, we found that there could still be microsaccade-related foveal SC motor bursts at the exact same time as visually-driven peripheral SC bursts (64), and we again argued that lateral inhibition would not explain these observations. It would be interesting in the future to understand the conditions under which physiological correlates of a lack of lateral inhibition would be most likely to occur.

Finally, the caution raised by our present work here is that in the classic study of saccade tasks, we may have generally assumed that the fixation spot was not influential. However, it clearly matters. While we believe that the foveal bursts that we observed are not direct responses to fixation spot removal in our tasks, it is still imperative to next ask whether they would continue to happen when there is no foveal sensory transient. For example, one could use a more abstract go instruction (that is also not represented foveally). One possibility could be to maintain the fixation spot visible throughout the trials, and instead use a subtle spatially-uninformative auditory cue as the instruction to generate a saccade. Under certain circumstances, such sounds only minimally affect visually-driven effects (81), and as long as they do not drive foveal SC neurons, one can record from the foveal SC and check whether the foveal bursts that we observed would still happen.

## Grants

We were funded by the German Research Association (Deutsche Forschungsgemeinschaft; DFG) under the Special Priority Programme “SPP 2411 Sensing LOOPS” (project numbers 520617944, 520283985; HA6749/11-1). We were also supported by the DFG-funded International Research Training Group IRTG2804.

## Author contributions

Conceived and designed research: TZ, AFD, ZMH

Analyzed data: TZ, AFD, ZMH

Interpreted results of experiments: TZ, AFD, ZMH

Prepared figures: TZ, ZMH

Drafted manuscript: TZ, AFD, ZMH

Edited and revised manuscript: TZ, AFD, ZMH

Approved final version: TZ, AFD, ZMH

